# Locomotor activity modulates associative learning in mouse cerebellum

**DOI:** 10.1101/099721

**Authors:** Catarina Albergaria, N. Tatiana Silva, Dominique Pritchett, Megan R. Carey

## Abstract

Changes in behavioral state are associated with modulation of sensory responses across visual, auditory and somatosensory cortices. Here we show that locomotor activity independently modulates performance in delay eyeblink conditioning, a cerebellum-dependent form of associative learning. Increased locomotor speed in head-fixed mice was associated with earlier onset of learning and trial-by-trial enhancement of learned responses. The influence of locomotion on conditioned responses was dissociable from changes in arousal and was independent of the sensory modality of the conditioned stimulus. Eyelid responses evoked by optogenetic stimulation of mossy fiber terminals within the cerebellar cortex, but not at sites downstream, were also positively modulated by ongoing locomotion. We conclude that locomotor activity modulates delay eyeblink conditioning through mechanisms acting on the mossy fiber pathway within the cerebellar cortex. Taken together, these results suggest a novel role for behavioral state modulation in associative learning and provide a potential mechanism through which engaging in movement can improve an individual’s ability to learn.

## Introduction

Changes in behavioral state can have profound influences on brain function. Recent studies have shown that locomotor activity and arousal influence both spontaneous activity and sensory-evoked responses in mouse sensory cortex (Ayaz et al., 2013; Bennett et al., 2013; Niell and Stryker, 2010; Reimer et al., 2014; Schneider et al., 2014; Vinck et al., 2015; Williamson et al., 2015; Zhou et al., 2014), with important consequences for sensory perception. To what extent the effects of locomotor activity generalize to other brain functions, however, remains unclear. Moreover, although locomotion also modulates activity within the cerebellar cortex (Ghosh et al., 2011; Hoogland et al., 2015; Ozden et al., 2012; Powell et al., 2015) the consequences of this modulation for cerebellar processing are largely unknown. Here we investigated the effects of locomotor activity on delay eyeblink conditioning, a cerebellum-dependent form of associative learning.

In delay eyeblink conditioning, animals learn to close their eye in response to an initially neutral conditioned stimulus (CS) that is reliably predictive of an aversive unconditioned stimulus (US), such as a puff of air to the eye (Gormezano et al., 1983; Kim and Thompson, 1997; Medina et al., 2000). CS and US signals are conveyed to the cerebellum via distinct input pathways, mossy fibers (CS) and climbing fibers (US) (De Zeeuw and Yeo, 2005; Yeo and Hesslow, 1998), each of which project to both the cerebellar cortex and deep cerebellar nucleus, where learning is thought to take place (Carey, 2011; Medina et al., 2000).

We find that locomotor activity improves associative learning through mechanisms that are dissociable from arousal and independent of the modulation of cortical sensory responses. Further, optogenetic circuit dissection revealed that locomotion modulates delay eyeblink conditioning through mechanisms acting on the mossy fiber (CS) pathway, within the cerebellar cortex.

## Results

We used a head-fixed apparatus with a freely rotating running wheel (Fig. 1A) (Chettih et al., 2011; Heiney et al., 2014b) to train mice in delay eyelid conditioning, a cerebellum-dependent form of associative learning (McCormick and Thompson, 1984; Steinmetz, 2000; Yeo and Hesslow, 1998). Daily conditioning sessions included 100 trials in which a neutral conditioned stimulus (CS, unless otherwise indicated, a white LED) was paired with an air puff unconditioned stimulus (US; a 50ms, 40psi air puff directed at the eye). The CS preceded the US by 300ms and the two stimuli co-terminated (Fig. 1B). Eyelid closures were recorded with a high-speed (900 fps) video camera and conditioned (CR) and unconditioned response (UR) amplitudes were extracted with offline image processing (Fig. 1B). We measured the mouse’s running continuously with an infrared sensor placed underneath the wheel (Fig. 1C).

**Fig. 1.**
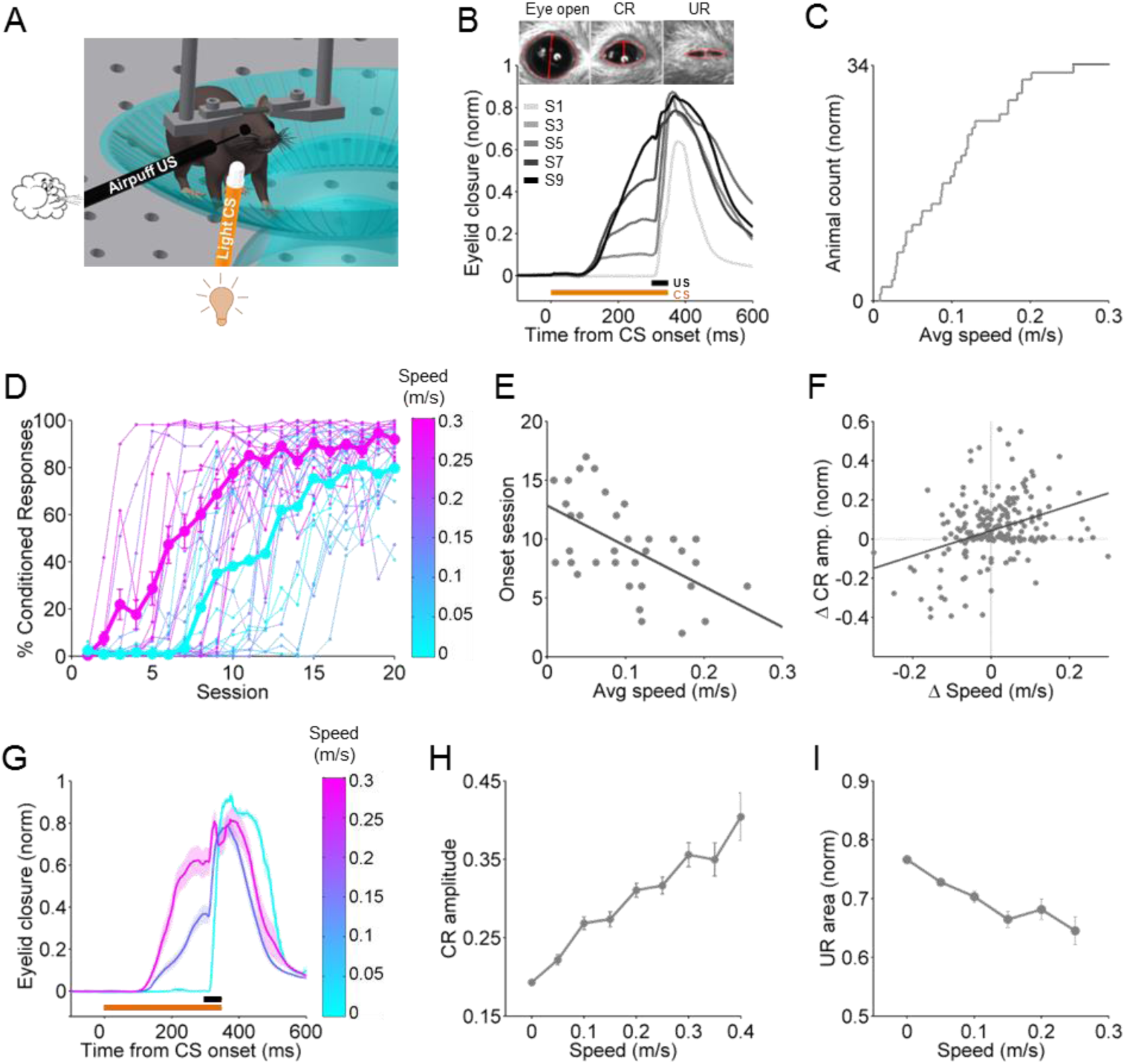
Eyeblink conditioning performance correlates with locomotor activity across mice, sessions, and trials. **A.** Setup schematic for eyeblink conditioning in head-fixed mice on a running wheel, illustrating a white LED as the CS and an air puff US. **B.** Average eyelid closure for a representative animal across 9 learning sessions (S1-S9). Each trace represents the average of 100 paired trials from a single session. Example video frames (acquired at 900 fps under infrared light) illustrate automated extraction of eyelid movement amplitude. **C.** Cumulative histogram of locomotor activity of all animals (n=34) trained with a light CS, calculated by summing the average speed of each session and dividing by the number of learning sessions (20). **D.** Learning curves for all animals, color-coded for their average running speed. Averages of the 20% fastest and slowest mice are superimposed in magenta and cyan, respectively. **E.** Onset session of learning for each animal, plotted against the animals’ average walking speed. Onset session was defined as the session in which the average CR amplitude exceeded 0.1. Each dot represents an animal. The line is a linear fit. See also Supp. Fig. 1. **F.** Each dot represents the relationship between session-to-session changes in locomotor activity (x-axis) and session-to-session changes in average CR amplitude (y-axis), for all sessions with significant change in the distribution of trials (compared to the previous session). The line is a linear fit. **G.** Average of trials from session 6 of one animal, divided into 3 speed intervals (0.05-0.1m/s; 0.2-0.25m/s; >0.35m/s) and color-coded accordingly. Shadows represent SEM. **H.** Correlation between CR amplitude and walking speed from trial-to-trial. CR amplitudes for all trials from the session where each individual animal crossed a threshold of 50% CR plus the following session are plotted. Speed was binned into 0.05 m/s speed bins. Error bars indicate SEM. **I.** Correlation between unconditioned response (UR) magnitude and walking speed from trial-to-trial. The normalized area under the UR in response to the air puff is plotted for Session 1, before emergence of conditioned responses. Speed was binned into 0.05 m/s speed bins. Error bars indicate SEM.

### Locomotor activity is associated with enhanced learning across mice

All mice trained with a light CS (n= 34) learned to reliably make well timed conditioned responses (CRs), steadily increasing both the percentage of trials that yielded conditioned responses and their amplitude over the course of the daily training sessions (Fig. 1B,D). However, the rate of learning was variable across individuals (Fig. 1D).

How much each animal ran on average across the 20 training sessions was also highly variable (Fig. 1C). Comparing acquisition curves between the mice that ran the most and those that ran the least revealed that on average, more active animals acquired conditioned responses more rapidly (Fig. 1D). For each animal, we subdivided the acquisition of CRs into three phases (Supp. Fig. 1) and analyzed: the onset session of learning (Fig. 1E), the slope of the acquisition curve (Supp. Fig. 1B), and the plateau value of CR amplitude (Supp. Fig. 1C). All of these features were positively correlated with the animals’ average running speeds (onset value: slope=-34.4, p=0.00057; slope value: slope=4.6, p=0.0017; plateau value: slope=0.85, p=0.038), indicating that the more an animal ran on average, the earlier, faster and better it learned.

### Conditioned response amplitudes correlate with locomotor activity across sessions and trials

Over the course of conditioning sessions the general trend is to increase frequency and amplitude of CRs until they asymptote. However, even in mice that reach high levels of performance, there can be variation in conditioned response amplitudes from session to session. We found that fluctuations in conditioned responses from one session to the next were positively correlated with changes in the amount of locomotor activity (Fig. 1F: slope=0.64, p<0.0001). Thus, learning is correlated with locomotor activity not just across animals, but also across sessions.

There is also considerable trial-to-trial variation in the amplitude of the conditioned responses within individual sessions (Fig. 1G,H). To determine whether this variation was correlated with trial-to-trial changes in locomotor activity, we sorted trials by running speed and averaged their conditioned response amplitudes. There was a linear positive relationship (one-way ANOVA on linear mixed effects model (LME), F(1,7202) = 352.52, p<0.0001) between running speed and conditioned response amplitude (Fig. 1H), indicating that locomotor activity and eyelid conditioning performance are correlated on a trial-by-trial basis.

We next asked whether the locomotor enhancement of CRs was specific to learned, conditioned responses, or if eyelid closures were generally affected, by analyzing the blinks (unconditioned responses, URs) that the animal made in response to the unconditioned air puff stimulus. We found that unconditioned response magnitudes were in fact negatively (one-way ANOVA on LME, F(1,1804) = 101.71, p<0.0001), not positively, modulated by locomotor speed (Fig. 1I). This indicates that locomotor enhancement is specific to conditioned responses, and further suggests that increased salience of the unconditioned response, for example through stronger climbing fiber input, does not account for the improved learning.

Taken together, these results indicate that there is a positive correlation between locomotor activity and eyelid conditioning performance, a relationship that holds true across animals, sessions and trials.

### Externally controlled changes in running speed are sufficient to modulate learning

When mice are free to initiate locomotor activity voluntarily, they run more during periods of high arousal (McGinley et al., 2015b). To ask whether the enhanced learning performance observed in Fig. 1 was driven indirectly by these changes in arousal, or by locomotor activity itself, we used a motorized treadmill to control running speed. We reasoned that if locomotor activity *per se* were driving the effect we observed with self-paced running, then running on a motorized treadmill should recapitulate it. We trained two randomly assigned groups of mice to a light CS at different fixed speeds, 0.12m/s (n= 5) or 0.18m/s (n=7), for the entire duration of each of the conditioning sessions. Mice were first habituated to the motorized wheel without the presentation of any stimuli until they walked normally and displayed no external signs of distress at the assigned speed (Supp. Movie 1).

Overall, mice learned very well while running on a motorized treadmill (Fig. 2A). Acquisition of CRs was generally faster and less variable than on the freely-rotating, self-paced treadmill (Fig. 2B: slope=-49.8; p<0.0001). Acquisition rate and conditioned response amplitudes were also running speed-dependent. Animals running at a faster speed learned earlier (Fig. 2A,B) and had larger conditioned response amplitudes (Fig. 2C) than mice on the slower treadmill.Interestingly, variability in learning was reduced on the motorized treadmill, likely due to the elimination of variability in running speed.

**Fig. 2.**
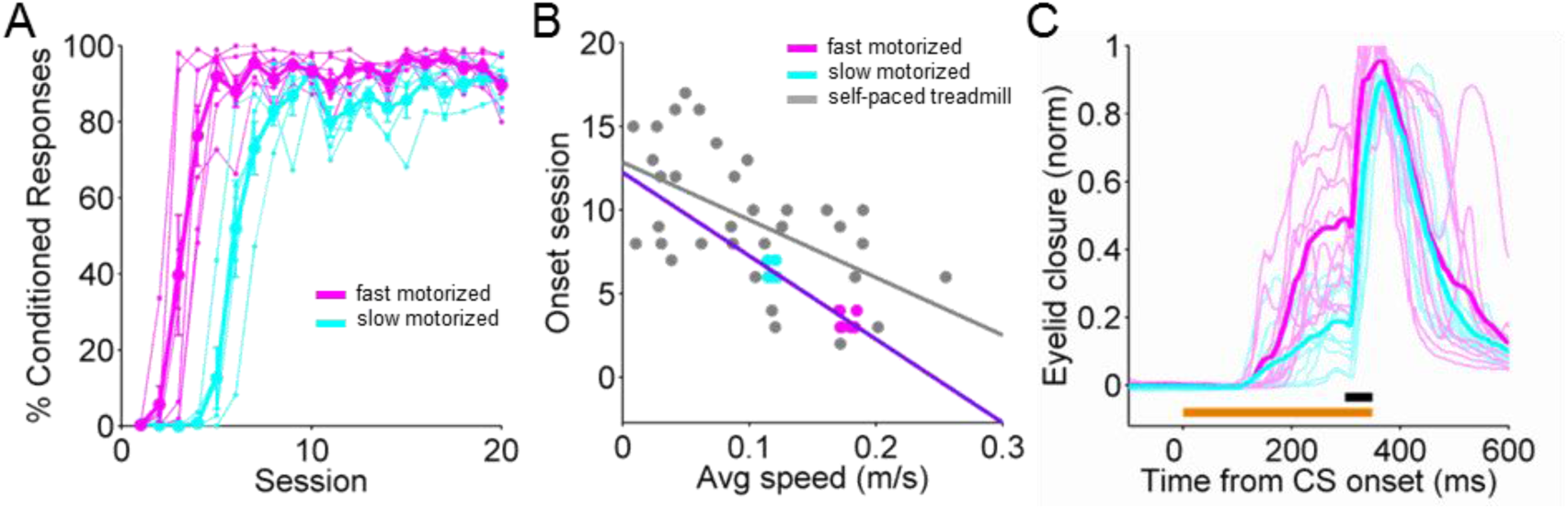
Speed-dependent modulation of eyeblink conditioning on a motorized treadmill. **A.** Individual learning curves with averages superimposed of two groups of mice, run on either a faster (magenta, 0.18m/s) or slower (cyan, 0.12m/s) motorized treadmill. **B.** Onset learning session for each animal. Fast (magenta) and slow (cyan) motorized data are superimposed on the self-paced treadmill data from Fig. 1E (gray). The purple line is a linear fit. **C.** Eyelid traces of individual trials for an animal on the fast (magenta) versus an animal on the slow (cyan) motorized treadmill at learning session 7. Only the traces for every 10th trial are shown.

The finding that externally imposed changes in running speed are sufficient to modulate learning extends the correlative data from the self-paced treadmill and suggests that locomotor activity *per se* modulates eyeblink conditioning performance in mice.

### Modulation is CS-independent and dissociable from changes in arousal

Behavioral state differentially affects processing of sensory stimuli of different modalities (Gentet et al., 2010; McGinley et al., 2015b). In particular, increased locomotion is associated with positive modulation of visual responses (Bennett et al., 2013; Chiappe et al., 2010; Keller et al., 2012; Niell and Stryker, 2010), but negatively modulates responses to auditory stimuli (Schneider et al., 2014; Zhou et al., 2014). We therefore wondered whether the correlation we observed between locomotor activity and learning performance could be due to enhanced sensory processing of our visual conditioned stimulus. To test this, we repeated the experiments from Fig. 1 but replaced the visual light CS with either a tone or a puff of air or vibratory stimulus delivered to the whisker (Fig. 3A). We found that learning to a whisker CS (n=25) proceeded marginally significantly (slope=-15.7; p=0.052) faster in mice that ran more rather than less (Fig. 3B). For the tone CS (n=16), we observed a similar, though not statistically significant, trend (Fig. 3B; slope=-8.9; p=0.49). From trial-to-trial, there was a positive correlation between running speed and conditioned response amplitude for both auditory (tone) and somatosensory (whisker) conditioned stimuli (Fig. 3C).

**Fig. 3.**
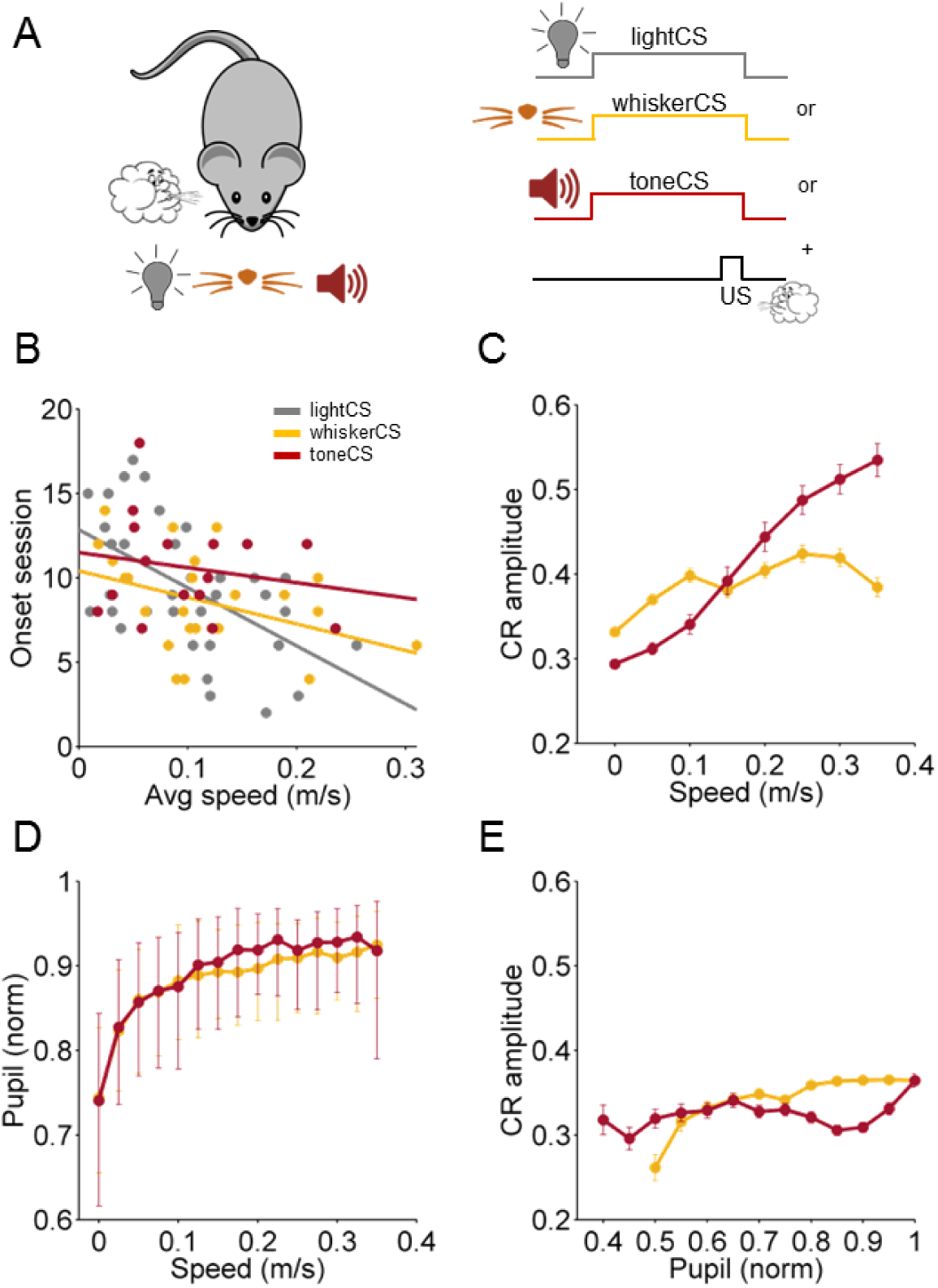
Modulation of CR acquisition and amplitude are CS-independent and dissociable from effects of arousal. **A.** Schematic for experiments using conditioned stimuli of different sensory modalities: light CS in gray, whisker CS in orange and tone CS in red; each was paired with an airpuff US. **B.** Onset learning session for animals from all three CS modalities, color-coded as in (A), plotted against the average running speed of each animal. **C.** Correlation between CR amplitude and walking speed for all trials with CRs from all training sessions using a whisker or tone CS. Data of trials from animals of each CS modality was separately averaged into 0.05 m/s speed bins. The lines plotted represent the average CR amplitudes against the average walking speed from trial-to-trial. Error bars indicate SEM. See Supp. Table 1 for GLM. **D.** Correlation between pupil size and walking speed for all trials. Median pupil sizes for each speed bin are plotted +/− quartiles (25th-75th percentile). Speed was binned into 0.025 m/s speed bins. **E.** Relationship between CR amplitude and pupil size for all CR trials from training sessions using a whisker CS (orange line) or a tone CS (red line). Data of trials from animals of each CS modality was separately averaged into 0.05 pupil size bins and the lines plotted represent the CR amplitude against the average pupil size from trial-to-trial. Error bars indicate SEM. See Supp. Table 1 for GLM.

Using a non-visual conditioned stimulus also allowed us to measure pupil diameter as a proxy for arousal (Joshi et al., 2016; Kloosterman et al., 2015; McGinley et al., 2015a; Reimer et al., 2014; Vinck et al., 2015). As has been previously shown, we found that on average, locomotor activity and pupil size were positively correlated with each other (Fig. 3D). Somewhat surprisingly, however, there was no obvious relationship between conditioned response amplitude and pupil size from trial-to-trial (Fig. 3E).

To investigate to what extent arousal vs. locomotor activity *per se* influenced conditioned response amplitudes from trial-to-trial, we fit the data with a Generalized Linear Model (GLM, Supp. Table 1). We found that for all CS modalities (visual, whisker and auditory), increased locomotor activity was a positive predictor of larger CR amplitude. Interestingly, larger pupil sizes were actually negative predictors of CR amplitude for auditory CS’s. This difference may be due to the previously observed differential influences of arousal on sensory processing of auditory vs. visual stimuli described above (McGinley et al., 2015b). The competition between the positive effects of locomotion and the negative effects of arousal revealed by the GLM for the auditory CS may also account for the weaker correlation between locomotor activity and acquisition rate in the tone CS experiments that we observed in Fig. 3B.

Thus, locomotor activity is associated with larger conditioned responses regardless of the sensory modality of the CS, and this effect can be dissociated from the (sometimes conflicting) effects of arousal.

### Locomotor activity modulates eyelid closures by acting downstream of mossy fiber inputs to the cerebellar cortex

The finding that locomotor activity positively modulated conditioned responses regardless of CS modality (Fig. 3; Supp. Table 1) suggested to us that locomotor activity might act downstream of CS processing, in eyelid conditioning areas themselves. In delay eyelid conditioning, the CS is conveyed to the cerebellum via mossy fiber inputs. Mossy fibers project both directly to the deep cerebellar nuclei and to the cerebellar cortex, where they synapse onto granule cells. Granule cell axons, the parallel fibers, project to Purkinje cells, which in turn inhibit deep nucleus neurons, converging once again with the direct mossy fiber-to-deep nucleus inputs (Fig. 4A).

**Fig. 4.**
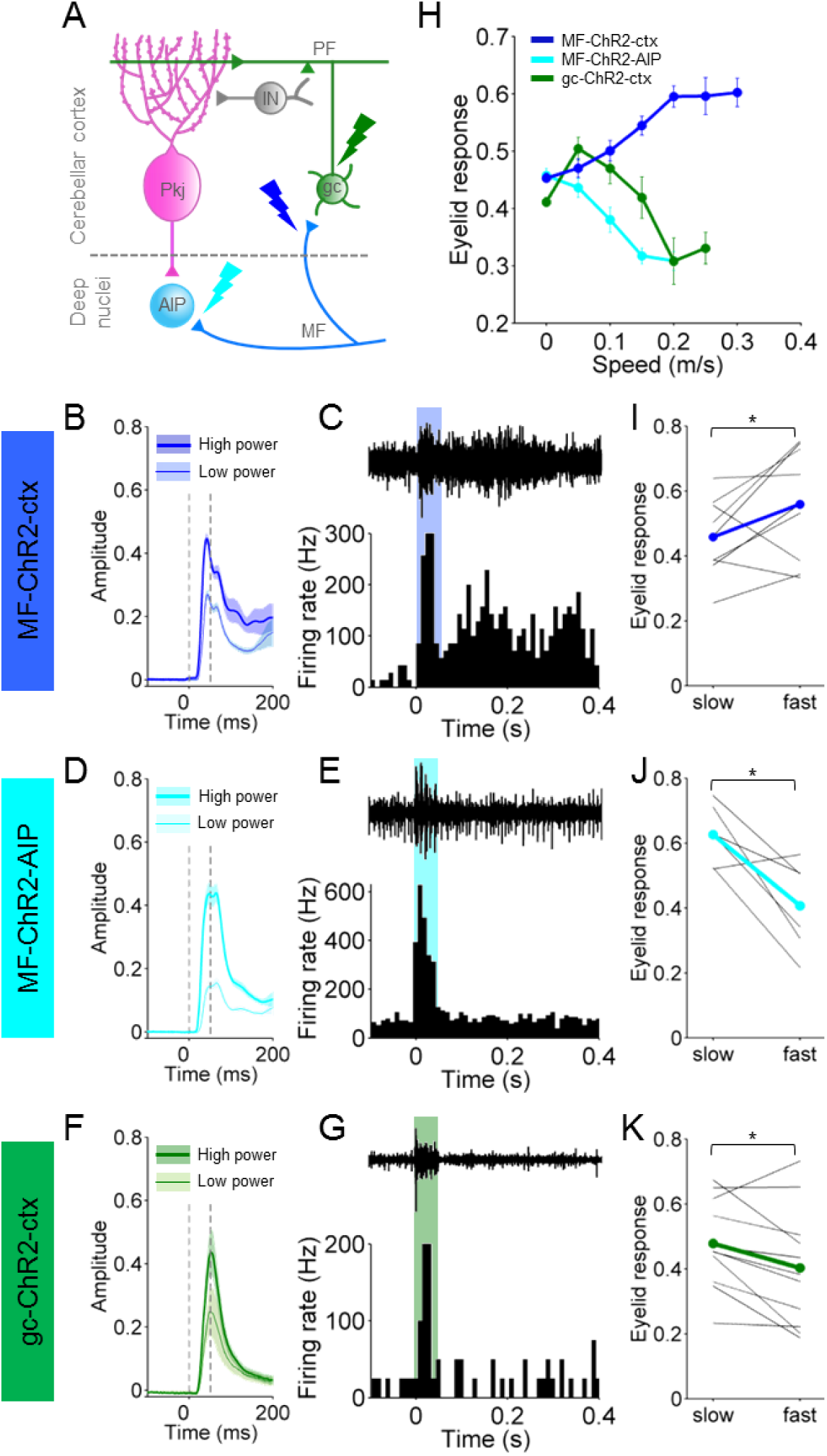
Eyelid closures evoked by optogenetic MF stimulation in the cerebellar cortex are positively modulated by locomotion. **A.** Schematic of the cerebellar circuit including some of the major cell types in the cerebellar cortex and deep nuclei. Lightning bolts represent the different targets of laser stimulation: mossy fibers (MF) terminals in the cerebellar cortex (blue), Mf terminals in the deep nuclei (cyan) and granule cells (gc, green). Pkj, Purkinje cell; IN, interneuron; PF, parallel fiber; AIP, anterior interpositus. **B D F** Eyelid movements in response to supra-threshold laser stimulation (473nm light pulses at 100Hz for 50ms) at two different intensities. Shadows indicate SEM. Laser pulse duration is indicated by the vertical dashed lines. An optical fiber was placed in an identified eyelid-related region of cerebellar cortex (**B**, blue) or AIP (**D**, cyan) of Thy1-ChR2-YFP mice that express ChR2 in cerebellar mossy fibers. In mice that express ChR2 in cerebellar granule cells (Gabra6cre-ChR2-YFP), an optical fiber was placed in the same eyelid region of cerebellar cortex (**F**, green). **C E G** *In vivo*, 1MOhm multiunit tetrode recordings in response to 50 ms light activation of ChR2 in awake mice. Example extracellular traces are shown above the peri-stimulus time histograms for each experimental condition. **C.** *In vivo* recordings from units in cerebellar cortex in response to MF-ChR2-ctx stimulation. Laser pulse duration is indicated by the blue shadow. **E.** *In vivo* recordings from units in cerebellar nuclei in response to MF-ChR2-AIP stimulation. Laser pulse duration is indicated by the cyan shadow. **G.** *In vivo* recordings from units in cerebellar cortex in response to gc-ChR2-ctx stimulation. Laser pulse duration is indicated by the green shadow. **H.** Correlation between laser-driven eyelid responses and walking speed from trial-to-trial. Speed was binned into 0.05 m/s speed bins. Error bars indicate sEm. See Supp. Table 1 for GLM. **I J K** Average amplitude of laser-elicited eye closures from animals walking at slow (0.06m/s) or fast (0.18m/s) pace, as set by the motorized treadmill. Each animal was tested for the two speeds within one session, using the same stimulation protocol and laser intensity. The average from all animals is superimposed.

To localize the modulation of eyelid closures by locomotor activity, we took advantage of this well-defined functional anatomy to directly optogenetically stimulate specific neurons within the mossy fiber CS pathway. We reasoned that optogenetic stimulation of neurons upstream, but not downstream, of the site of locomotor modulation would evoke eyelid closures that would be positively modulated by locomotor activity.

We therefore implanted fiber optics into the cerebellar cortex or deep nucleus of Thy1-ChR2/EYFP transgenic mice expressing channelrhodopsin 2 (ChR2) in cerebellar mossy fibers (from here on termed MF-ChR2 mice; Supp. Fig. 2A, B) (Arenkiel et al., 2007; Hull and Regehr, 2012). As expected (Heiney et al., 2014a; Steinmetz and Freeman, 2014), optogenetic stimulation in the eyelid region of cerebellar cortex (Fig. 4B,C) or the anterior interpositus deep cerebellar nucleus (AIP, Fig. 4D,E) of MF-ChR2 mice evoked increases in multiunit neural activity (Fig. 4C,E) and short-latency, stimulation-intensity dependent eyelid closures (Fig. 4B,D).

We found that while running on a self-paced treadmill, eyelid closures evoked by optogenetic stimulation of MFs within the cerebellar cortex (MF-ChR2-ctx, n=9) were larger on trials in which mice were running faster (Fig. 4H, blue). Interestingly, this effect was highly location specific. In the same mouse line, optogenetic stimulation of MF terminals in the AIP (MF-ChR2-AIP, n=5) evoked eyelid closures that were not positively correlated with locomotor speed (Fig. 4H, cyan). Thus, the effects of MF-ChR2 stimulation in the cerebellar cortex vs. deep nucleus were differentially (see Supp. Table 1 for GLM results) modulated by locomotion. These results suggest that locomotor activity acts downstream of MF inputs to the cerebellar cortex to modulate eyelid closures.

Mossy fibers in the cerebellar cortex synapse directly onto cerebellar granule cells. To investigate whether the locomotor modulation of eyeblinks was occurring downstream of granule cells, we analyzed eyelid closures in response to optogenetic stimulation in mice expressing ChR2 in cerebellar granule cells (from here on termed gc-ChR2 mice; Supp. Fig. 2C) (Fünfschilling and Reichardt, 2002; Madisen et al., 2012).

As with MF stimulation, optogenetic granule cell stimulation within the eyelid area (gc-ChR2-ctx, n=11) evoked stimulus intensity-dependent short-latency eyelid closures and changes in multiunit activity (Fig. 4F,G). However, on a self-paced treadmill, eyelid closures evoked by optogenetic stimulation of granule cells within the cerebellar cortex (Fig. 4H, green) were not positively correlated with locomotor activity (see Supp. Table 1 for GLM results).

The same pattern of dependence on locomotor activity was observed in mice running on a motorized treadmill. Again, MF-ChR2-ctx (n=7) eyelid closures were larger (p=0.03; paired t-test) when the animals were running on a fast (0.18 m/s) vs. slow (0.06 m/s) motorized treadmill (Fig. 4I-K). However, MF-ChR2-AIP (n=6) and gc-ChR2-ctx (n=11) responses were suppressed (p=0.03, p=0.0197; paired t-test). This result is reminiscent of the suppression of unconditioned responses (Fig. 1I), suggesting that there may be a generalized suppression within the eyelid closure pathway, likely downstream of mossy fibers.

Thus, while optogenetic stimulation of granule cells and MFs in either cerebellar cortex or deep nucleus readily evoked eyelid closures, only cortical mossy fiber stimulation yielded responses that were positively correlated with locomotion. This remarkable location and cell-type specificity suggests that locomotor activity exerts its influence within the cerebellar cortex, possibly at the MF-granule cell connection itself.

Interestingly, in the naïve state, mossy fiber responses to a sensory conditioned stimulus are not sufficient to drive eyelid closures – only learned, conditioned responses pass through the cerebellar cortex. Therefore, the localization of the locomotor enhancement of eyelid closures to the cerebellar cortical MF pathway suggests that the modulation may be a particular feature of learned eyelid closures, an idea that we test with the next experiment.

### Learning to an optogenetic conditioned stimulus is positively modulated by locomotor activity

Our results so far have suggested that 1) locomotor activity modulates eyeblink conditioning (Figs. 1-3), and 2) locomotor activity modulates eyelid closures evoked via MF projections to the cerebellar cortex (Fig. 4). These findings raised the intriguing possibility that locomotor activity also modulates learning within the cerebellar cortex. If so, then we should observe an earlier onset of learning and trial-to-trial modulation of conditioned responses if optogenetic stimulation of cerebellar cortical MFs were used as the conditioned stimulus. Although optogenetic mossy fiber stimulation has not previously been shown to act as a conditioned stimulus for eyeblink conditioning, electrical stimulation of MFs can substitute for a sensory CS to drive eyelid conditioning in rabbits (Kreider and Mauk, 2010; Lavond et al., 1987; Steinmetz et al., 1986).

Therefore, we asked 1) whether optogenetic stimulation of MF terminals within the cerebellar cortex could act as a conditioned stimulus for eyeblink conditioning (Fig. 5A,B), and 2) if so, if the elicited conditioned responses would be modulated by locomotor activity. We placed an optical fiber in the eyelid region of the cerebellar cortex, as above, but reduced the stimulation intensity so that no eyelid movement was evoked in response to CS stimulation (Fig. 5C).

**Fig. 5.**
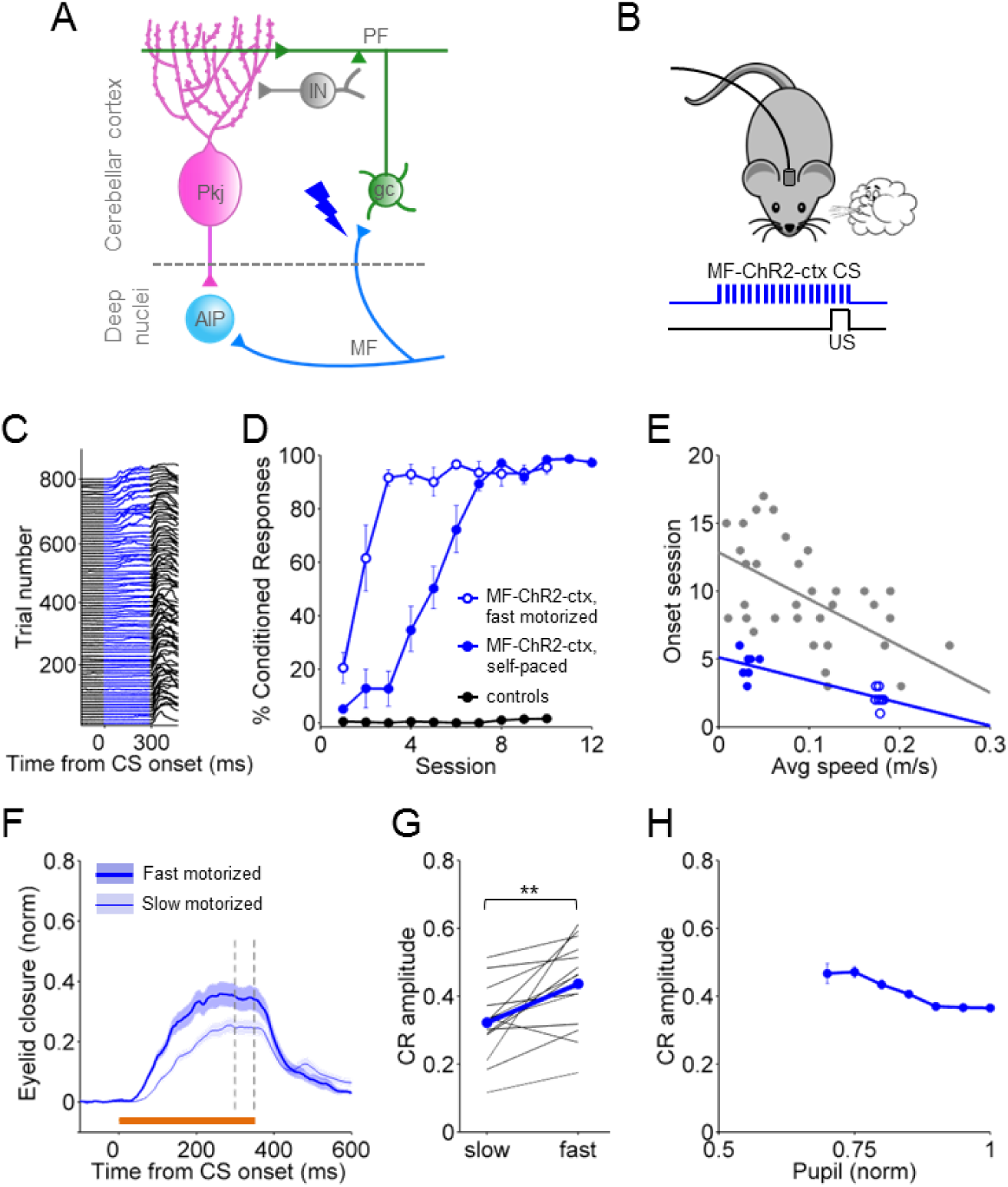
Conditioned responses acquired with optogenetic stimulation of cerebellar mossy fibers in the cerebellar cortex are positively modulated by locomotor activity. **A.** Cerebellar circuit diagram with a blue lightning bolt representing the site of laser stimulation: mossy fiber terminals in eyelid-related cerebellar cortex. **B.** Schematic of eyelid conditioning protocol using MF-ChR2-ctx optogenetic stimulation as a replacement for the CS. Animals were trained to a sub-threshold (i.e. not eliciting eye movement) laser stimulation of mossy fibers (473nm light pulses at 100Hz for 350ms) paired with an airpuff to the eye as US. **C.** Representative individual trials for one mouse trained with a MF-ChR2-ctx CS and airpuff US, during the first eight sessions. The eyelid trace for every 9th trial is plotted. **D.** %CR Learning curves to MF-ChR2-ctx CS's for animals walking on a self-paced treadmill (filled circles), and animals running at a fast fixed speed (0.18m/s) on the motorized treadmill (open circles). To control for the possibility that the laser stimulation was inadvertently acting as a visual CS, wildtype controls not expressing ChR2 were implanted with optical fibers and underwent the same training protocol (black circles). **E.** Onset learning session for each animal from the self-paced treadmill group (filled blue circles) and the fast motorized treadmill group (open blue circles). are superimposed on the self-paced treadmill data from Fig. 1E (gray). The blue line is a linear fit. **F G H** After learning reached a plateau, both groups were tested for the expression of CRs at two fixed speeds set on the motorized treadmill: slow (0.06m/s) and fast (0.18m/s). **F.** Average of CS-only trials from the slow (thin line) and fast (thick line) blocks of trials, for one representative animal. Shadows indicate SEM. **G.** Average CR amplitude of responses from each animal walking at slow (0.06m/s) vs fast (0.18m/s) pace on the motorized treadmill. The average from all animals is superimposed. **H.** Relationship between CR amplitude and pupil size from trial-to-trial on the motorized treadmill at the two tested speeds. Error bars indicate SEM.

As with a sensory CS, mice gradually learned to make well-timed eyelid closures in response to optogenetic stimulation of cortical MFs (Fig. 5C). On a self-paced treadmill, animals (n=7) acquired CRs at rates that were, overall, comparable to those conditioned with a sensory CS (Fig. 5D, blue filled circles). However, given that these particular mice exhibited low levels of spontaneous locomotor activity, their acquisition was faster than would have been expected based on the relationship between locomotion and learning onset we described in Figs. 1 and 2 (Fig. 5E).

We next asked whether animals would acquire conditioned responses to a cortical MF-optogenetic CS even faster if they were running on a motorized treadmill. Because learning was already quite rapid, we were surprised to find that running on a fast motorized treadmill (n=7) accelerated the onset of learning to a mossy fiber conditioned stimulus (Fig. 5 D,E: slope=-16.8; p=0.00015).

To determine whether locomotor activity modulated the expression of conditioned responses, animals (n=14) were placed on the motorized wheel for post-conditioning test sessions with 50% CS+US and 50% CS-only trials (Chettih et al., 2011). Each animal was tested at two speeds (0.06m/s and 0.18m/s; trials were presented in speed blocks and the order of blocks was counterbalanced across animals) (Fig. 5F,G). CR amplitudes were larger when the animals were running at faster speeds (Fig. 5F,G: p=0.0017; paired t-test). Finally, the locomotor modulation of optogenetically evoked CRs could not be accounted for by changes in arousal, as CRs were not significantly correlated with pupil size (Fig. 5H: slope=-0.39; p=0.15; GLM).

Thus, locomotor activity improves learning to a conditioned stimulus consisting of direct optogenetic stimulation of mossy fibers within the cerebellar cortex.

## Discussion

We found that delay eyeblink conditioning is modulated by locomotor activity in mice. Both acquisition and expression of learning was enhanced in a speed-dependent manner when animals were running on either a self-paced or motorized treadmill. The effects of locomotor activity were specific to learned responses and were dissociable from changes in arousal. Modulation of eyelid responses by arousal, like the modulation of cortical sensory responses, varied across sensory modalities. In contrast, locomotor activity consistently enhanced performance across CS modalities. Bypassing sensory processing by direct optogenetic activation of mossy fiber projections revealed that the effects of locomotor activity acted downstream of these inputs, within the cerebellar cortex. Our findings suggest a novel role for behavioral state in the modulation of associative learning and suggest that an individual’s ability to learn is influenced by ongoing movement and may be improved by engaging in motor activities.

### Dissociable effects of locomotion and arousal

Several lines of evidence suggest that the positive correlation we observe between locomotor activity and learning is distinct from previously described (McGinley et al., 2015a; Vinck et al., 2015) modulation of cortical sensory responses. First, while self-paced running speed is influenced by arousal, in the case of the motorized treadmill, the speed was externally imposed. Nevertheless, responses were locomotor speed-dependent regardless of whether running was self-paced or on a motorized treadmill. Second, in experiments with auditory and whisker conditioned stimuli, a generalized linear model revealed independent contributions of pupil size and locomotor speed. In particular, responses to an auditory CS were negatively modulated by arousal, consistent with previous findings that have shown differential effects of arousal on auditory vs visual cortical responses (McGinley et al., 2015a; McGinley et al., 2015b; Niell and Stryker, 2010; Schneider et al., 2014; Zhou et al., 2014). Along those same lines, bypassing noise in sensory processing could explain why mice learned faster to an optogenetic mossy fiber CS. In general, the outcome of learning is likely to depend on both the quality of the sensory representation of the CS as conveyed to the cerebellum via mossy fibers, and the locomotor-dependent modulation of cerebellar processing that we describe here.

### Locomotor activity modulates eyeblink conditioning within the cerebellar cortex

Several previous studies have highlighted the difficulties in establishing robust eyeblink conditioning in head-fixed mice (Boele et al., 2009; Koekkoek et al., 2002). Performance improved when mice were allowed to run freely on a running wheel, an observation that has been attributed to removing stress and allowing the animals to engage in one of their favorite activities (Chettih et al., 2011; Heiney et al., 2014b). Our finding that locomotor activity modulates delay eyelid conditioning directly, within the cerebellar cortex, provides a possible alternative mechanism to account for the earlier difficulties and the success of more recent approaches.

Eyelid closures evoked by optogenetic stimulation of mossy fiber terminals within the cerebellar cortex, but not in the AIP or with granule cell stimulation, were positively modulated by increased locomotor speed. Moreover, when optogenetic stimulation of cerebellar cortical mossy fibers was used as a conditioned stimulus, learning was accelerated and there was trial-to-trial enhancement of learned responses. These effects were also speed dependent and independent of changes in arousal. Taken together, these results suggest that locomotor activity modulates eyeblink conditioning by acting on the mossy fiber pathway, within the cerebellar cortex itself.

We found that optogenetic stimulation of mossy fiber terminals within the cerebellar cortex, but not stimulation of their post-synaptic targets, cerebellar granule cells, evoked eyelid responses that were positively modulated by locomotion. This surprising finding suggests that the mechanism of action of locomotor activity is highly localized to the MF-granule cell connection.

The cerebellum receives many neuromodulatory inputs (Ross et al., 1990; Schweighofer et al., 2004), in particular, noradrenergic signals relating to arousal (Aston-Jones and Cohen, 2005; Reimer et al., 2014) that are known to affect cerebellar circuit activity in a variety of ways (Bloom et al., 1971; Carey and Regehr, 2009; Dieudonné, 2001; Olson and Fuxe, 1971; Paukert et al., 2014). One possibility is that neuromodulator release related to locomotor activity positively modulates the effectiveness of MF-gc transmission by affecting some combination of mossy fiber terminals, granule cells, or Golgi cells. However, the ability to clearly dissociate effects of arousal from those of locomotor activity *per se* makes it unlikely that the effect we describe can be explained by a simple relationship between noradrenergic activation and performance (Joshi et al., 2016; Martins and Froemke, 2015).

Another intriguing possibility is that convergence of locomotor and CS signals onto individual granule cells allows them to convey more reliable CS information to interneurons and Purkinje cells (Bengtsson and Jörntell, 2009; Ishikawa et al., 2015; Ozden et al., 2012; Powell et al., 2015). Granule cells receive input from 3-5 distinct mossy fibers. Recent studies have demonstrated multimodal convergence of mossy fiber inputs onto single granule cells (Huang et al., 2013; Ishikawa et al., 2015; Sawtell, 2010). Granule cells require multiple simultaneous MF inputs to reach threshold (Jörntell and Ekerot, 2006), and MF-gc synaptic transmission is enhanced by glutamate spillover during locomotion (Powell et al., 2015). Convergence of locomotor and conditioned stimulus MF inputs to a granule cell would therefore be expected to increase the probability of a postsynaptic action potential in response to the CS during locomotion. Thus, it may not be necessary to invoke neuromodulatory mechanisms to explain the effects of locomotor activity on cerebellar circuit processing that we describe here.

## Methods

### Animals

All procedures were carried out in accordance with the European Union Directive 86/609/EEC and approved by the Champalimaud Centre for the Unknown Ethics Committee and the Portuguese Direcção Geral de Veterinária (Ref. No. 0421/000/000/2015). All procedures were performed in C57BL/6 mice approximately 10–14 weeks of age. The Thy1-ChR2/EYFP mouse line (Arenkiel et al., 2007) was obtained from The Jackson Laboratory (stock number: 007612). The Gabra6cre-ChR2-YFP mouse line was obtained by crossing Gabra6cre mice (Fünfschilling and Reichardt, 2002) with ChR2-EYFP-LoxP (Madisen et al., 2012) from The Jackson Laboratory (stock number: 012569). Male and female mice were housed in groups of 3-5 with food and water *ad libitum* and were kept on a reverse light cycle (12:12 hour light/dark) so that all experiments were performed during the dark period while mice were more active.

### Surgical procedures

In all our surgeries, animals were anesthetized with isoflurane (4% induction and 0.5 – 1% for maintenance), placed in a stereotaxic frame (David Kopf Instruments, Tujunga, CA) and a head plate was glued to the skull with dental cement (Super Bond – C&B). For *in vivo* electrophysiological recordings, craniotomies were drilled over the eyelid area of cerebellar cortex (RC -5.7, ML +1.9) and then filled over with a silicon based elastomer (Kwik-cast, WPI) that was easily removed just before recording sessions. For optogenetic manipulations, optical fibers with 200μm core diameter, 0.22 NA (Doric lenses, Quebec, Canada) were lowered into the brain through smaller craniotomies, and positioned at the cortical eyelid region (RC -5.7, ML +1.9, DV 1.5) or just over the AIP (RC -6, ML +1.7, DV 2.1) (Ohmae and Medina, 2015). The implants were fixed into place using dental cement (Super Bond – C&B). After any surgical procedure, mice were monitored and allowed ~1-2 days of recovery.

### Behavioral procedures

The experimental setup was based on previous work by (Chettih et al., 2011; Heiney et al., 2014b). For all behavioral experiments, mice were head-fixed but could walk freely on a Fast-Trac Activity Wheel (Bio-Serv) and habituated to the behavioral setup for at least 4 days prior to training. For experiments on the motorized treadmill, mice were additionally habituated to walk at the target speed until they walked normally and displayed no external signs of distress. To externally control the speed of the treadmill, a DC motor with an encoder (Maxon) was used. Running speed was measured using an infra-red reflective sensor placed underneath the treadmill. Eyelid movements of the right eye were recorded using a high-speed monochromatic camera (Genie HM640, Dalsa) to monitor a 172 × 160 pixel region, at 900fps. Custom-written software using LabVIEW, together with a NI PCIE-8235 frame grabber and a NI-DAQmx board (National Instruments), was used to trigger and control all the hardware in a synchronized manner.

After habituation, most of the behavioral experiments consisted of 3 phases: acquisition, test and extinction. Acquisition sessions consisted of the presentation of 90% CS-US paired trials and 10% CS-only trials, which allow for the analysis of the kinematics of CRs without the masking effect that comes from the US-elicited reflex blink. Test sessions were presented after mice had asymptote and consisted of 50% paired and 50% CS-only trials (Chettih et al., 2011). During extinction 100% of the trials presented were CS-only. Each session consisted of 110 trials, separated by a randomized inter trial interval (ITI) of 5-20s. In each trial, CS and US onsets were separated by a fixed interval (ISI) of 300ms and both stimuli co-terminated.

For all training experiments, the unconditioned stimulus (US) was an air puff (30-50psi, 50ms) controlled by a Picospritzer (Parker) and delivered via a 27G needle positioned ~0.5cm away from the cornea of the right eye of the mouse. The direction of the air puff was adjusted for each session of each mouse so that the unconditioned stimulus elicited a normal eye blink. The CS had a 350ms duration and was either a 1) white light LED positioned ~3cm directly in front of the mouse; 2) a piezoelectric device placed ~0.5cm away from the ipsilateral vibrissal pad; or 3) a tone (10kHz, 68dB) delivered by a speaker placed ~15cm away from the mouse.

### Behavioral analysis

The movie from each trial was analyzed offline with custom-written software using MATLAB (MathWorks). The distance between eyelids was calculated frame by frame by thresholding the gray scale image of the eye and extracting the count of pixels that constitute the minor axis of the elliptical shape that delineates the eye. Eyelid traces were normalized for each session, ranging from 1 (full blink) to 0 (eye fully open). Trials were classified as CRs if the eyelid closure reached at least 0.1 (in normalized pixel values) and occurred between 100ms after the time of CS onset and the onset of US. The average running for each animal was calculated by summing the average speed of each session (total distance run divided by session duration) and dividing by the total number of learning sessions, usually 20. Running speed for trial was calculated by dividing the distance run in the intertrial interval preceding the current trial by the elapsed time.

### Optogenetic stimulation

For driving a blink, either through the activation of ChR2 in mossy fibers (cerebellar cortex or AIP), or the activation of ChR2 in granule cells, a 50ms stimulation at 100Hz, 40% duty cycle was delivered with an intensity of 2 to 119 mW/mm2. Light from a 473 nm laser (LRS-0473-PFF-00800-03, Laserglow Technologies) was controlled with custom-written code using LabView software, and laser power was adjusted for each mouse and controlled for each experiment using a powermeter (Thorlabs) at the beginning and end of each session. To investigate the modulation of locomotion on laser-driven blinks, the intensity of the laser was adjusted so that it would elicit an intermediate eyelid closure.

For activation of ChR2 in mossy fibers as a CS, laser power was lowered until no eye movement was detected. A 350ms stimulation at 100Hz, 20% duty cycle was paired with the airpuff US. Power output per unit area ranged from 4 to 16mW/mm2.

### Electrophysiological recordings

Single and multi-unit recordings were performed with quartz-insulated tungsten tetrodes (Thomas Recording, tip type A, impedances between 1-3 MOhm) or Tungsten microelectrodes (75 um shaft diameter, FHC) mounted on a 200um diameter optic fiber (fiber placed with 300um offset to minimize the opto-electric artifact). Electrodes were controlled using a 3-axis stereotaxic manipulator (Kopf) or a motorized 4-axis micromanipulator (PatchStar, Scientifica).

Recordings were performed with an Intan digital amplifier/headstage with the Open Ephys digital acquisition board (Siegle et al., 2015). Online monitoring was done using a custom Bonsai software interface (Lopes et al., 2015). All recordings were digitized from the wide-band signal (0.1 Hz - 10kHz, sampled at 30kHz), and sorted offline using custom Matlab code for unit analysis. Single units were determined by checking the unit parameters, e.g. ISI, firing rate, and LCV (Van Dijck et al., 2013).

In awake, head-fixed mice running on a treadmill, we targeted the eyeblink microzone either by monitoring the local field potential for a large depolarization to an air puff stimulus applied to the ipsilateral eyelid (Ohmae and Medina, 2015) or by optogenetic stimulation driving a blink response.

We recorded from a total of 23 recording sites, from 6 animals (5 Thy1-ChR2/EYFP animals, 4 sites in AIP and 14 sites in the cerebellar cortex, and 1 Gabra6-ChR2 mouse with 5 recording sites). Recording sites were chosen for clear unit activity. However, across all recording sites, even in the presence of a well-isolated single unit, spike waveform and ISI changes during laser stimulation indicated the presence of multi-unit activity to optogenetic manipulation.

### Histology

After photostimulation experiments using chronically implanted optical fibers dipped in DiI (Sigma), animals were perfused transcardially with 4% paraformaldehyde and their brains removed, so that fiber placement could be examined (Supp. Fig. 2). Coronal sections were cut in a vibratome and mounted on glass slides with mowiol mounting medium. Histology images were acquired with an upright confocal laser point-scanning microscope (Zeiss LSM 710), using a 10x objective.

### Statistical analysis

All statistical analyses were performed using the Statistics toolbox in MATLAB. For the correlation between speed vs onset session (Fig. 1E; Fig. 2B; Fig. 3B; Fig. 5E) and speed vs slope or plateau (Supp. Fig. 1B, C), we used linear regression analysis. We also fitted a linear regression to delta speed vs delta CR amplitude (Fig. 1F). For the correlation between speed vs CR amplitude (Fig. 1H), and speed vs UR area (Fig. 1I), we used a one-way ANOVA applied to a linear mixed effects model (LME). To compare the effects of speed and arousal (pupil) on CRs (Fig. 3C, E) or on laser-driven eyelid responses (Fig. 4H), we used a generalized linear model (GLM). We used the GLM also to analyze the effect of arousal on CR amplitude when the speed was fixed by the motorized treadmill (Fig. 5H). To compare the amplitudes of eye closure on the fast and slow speeds on the motorized treadmill (Fig. 4I,J,K; Fig. 5G), we used a paired t-test. All t-tests were two-sided. Differences were considered significant at *p < 0.05, **p < 0.01 and ***p < 0.001. No randomization was used, but mice were assigned to specific experimental group without bias and no animals were excluded.

## Acknowledgements

We thank Tracy Pritchett for technical assistance and Patrícia Francisco for her help on some of the auditory CS experiments. We thank Gil Costa for illustrations and the Champalimaud Research Hardware Platform for technical support. We are grateful to the Carey lab for helpful discussions throughout the project and in particular to Hugo Marques and Jovin Jacobs for comments on the manuscript. We thank João Fayad and Michael Orger for advice on data analysis. This work was supported by grants from HHMI, Bial, FCT, and ERC.

